# Low determinism in pelagic fungal community assembly across climate zones in Scandinavian lakes

**DOI:** 10.1101/2024.06.04.597408

**Authors:** Pedro M. Martin-Sanchez, Christian Wurzbacher, Laurent Fontaine, Maria Jimenez Lara, Heli Juottonen, Nicolas Valiente, Even Werner, R. Henrik Nilsson, Alexander Eiler

## Abstract

In lakes, fungi play pivotal roles in biogeochemical processes, particularly in the decomposition of organic matter. However, these organisms have been historically underrepresented and taxonomically unresolved in previous research. In this study, we employ high-resolution sequencing to delve into the fungal diversity within two distinct Scandinavian lake datasets spanning a vast latitudinal gradient, diverse climatic zones ranging from nemoral to arctic, and a wide spectrum of nutrient conditions. Utilizing eukaryotic primers, we reveal that fungi contribute, on average, 2.07% in the Norwegian and 5.4% in the Swedish sequencing dataset, with remarkable spikes of up to 55% in forested lakes. Fungal-specific sequencing identified the prevalence of several fungal phyla, including *Ascomycota*, *Rozellomycota*, *Basidiomycota*, *Chytridiomycota*, *Aphelidiomycota*, and *Mortierellomycota*. Notably, our research uncovers a striking degree of variability in fungal communities across the studied lakes, defying correlations with measured environmental factors or geographic distance. Null model analyses suggest that deterministic processes do not consistently override the influence of ecological drift in shaping these biogeographic patterns. However, signals for dispersal limitations and mass effects could be detected. These findings challenge conventional paradigms of eukaryotic communities being strongly guided by deterministic processes and underscore the complexity of fungal community assembly processes across climate zones.

## INTRODUCTION

Fungi are abundant in many if not all aquatic ecosystems, and have been suggested to play an important role in biogeochemical cycles ^1–6^. They are key players in organic matter (OM) decomposition from freshwater environments, where fungi can comprise high relative abundances, constituting up to 50% of eukaryotic sequences ^7,8^. Nonetheless, our understanding of fungal diversity, quantitative abundance, ecological function, and, in particular, their community assembly and interactions with other microorganisms remain predominantly speculative, unexplored, and absent from contemporary concepts in aquatic ecology and biogeochemistry ^3–5,9,10^. Pioneering efforts have unveiled a high diversity of uncultivated fungal taxa, detected solely through molecular tools ^11,12^. They are termed dark fungal taxa as these lake and river fungi are uncharacterized ^13–15^.

In contrast to studies on soil ecosystems ^16,17^, studies on aquatic fungi are still insufficient or lack adequate spatial and temporal resolution to infer the natural history of freshwater fungi. An initial global survey compiled from the literature has provided valuable insights into the ecology and biogeography of aquatic hyphomycetes ^18^. Nonetheless, unraveling the ecology and biogeography of diverse fungal groups necessitates an extensive array of such studies, with sufficient environmental metadata, for shedding light on the dark fungal taxa and their role in aquatic ecosystems. Moreover, the mechanisms governing community assembly and the regulation of fungal diversity in aquatic ecosystems remain largely enigmatic. While prokaryotic community assembly is predominantly deterministic and to a large extent driven by environmental sorting ^19,20^, this may differ significantly in freshwater fungi where stochastic processes, dispersal limitation, biotic interactions, or historical processes may exert greater influence over community assembly.

To shed light on the ecological phenomena of community assembly, a statistical “toolbox” is used consisting of several complementary approaches that all have their strengths and limitations. Recently, null model approaches that facilitate quantitative comparisons of assembly processes have been increasingly used ^21,22^. Among these approaches, methods to discern the elements of metacommunity structure (EMS) have sought to distinguish between communities randomly assembled from those shaped by species-sorting processes ^23,24^. Co-occurrence indices ^25^ and incidence-based (Raup-Crick) beta-diversity (β_RC_) ^26^ have been used to parse deterministic and stochastic assembly processes ^27,28^. Additionally, the integration of phylogenetic information into null model approaches, based on the premise that phylogenetic relatedness mirrors shared environmental response traits ^29^, has expanded the toolkit on disentangling community assembly ^30^. Stegen *et al*. ^22,31^ have combined null model approaches based on phylogenetic and abundance-based (Raup-Crick) beta-diversity (β_RCbray_) measures. This approach enables the quantitative estimation of the relative importance of processes such as selection, drift, dispersal limitation, and mass effects, while also distinguishing between heterogenous (i.e., beta-diversity enhancing) and homogeneous (i.e., beta-diversity diminishing) selection processes.

In the present study, we use high-throughput amplicon sequencing of (i) a universal region of the small subunit (SSU) 18S rRNA gene (V6-V8) spanning all eukaryotic diversity, and (ii) other regions of the rRNA operon including the 5.8S gene and the internal transcribed spacer 2 (ITS2), which is recognized as a universal fungal barcode ^16,32^. Our main goal is to delve deeply into fungal diversity within two independent investigations of lakes across Scandinavia (Norway and Sweden), covering the nemoral to arctic zones. Through statistical analyses, we aim to unveil the environmental factors that most strongly relate to fungal diversity, including catchment land use/cover, hydrological properties, and geochemical parameters. We seek to disentangle the roles of random (i.e. ecological drift) and deterministic (i.e. environmental sorting and dispersal) factors in shaping the assembly of fungal communities across lake ecosystems.

## RESULTS and DISCUSSION

### Characteristics of surveyed Norwegian and Swedish lakes

In our study, we set out to investigate and quantify the diversity of epilimnic freshwater fungi across a broad range of latitudes, spanning from 55.4° to 68.3° (Figure S1). This represents globally the latitudes with the highest concentration, area, and perimeter of inland water bodies ^33^. The sampled Norwegian and Swedish lakes presented a diverse array of ecological contexts, covering vegetation zones from nemoral to arctic (subalpine). Metadata associated with each of the 160 sampled lakes, or at least a substantial subset thereof (61 Norwegian and 83 Swedish lakes where sufficient fungal sequences were retrieved), included latitude and longitude, temperature, chlorophyll concentration, nutrients, and catchment properties. Summary statistics of metadata are shown in the supplementary Table S1.

Beyond the distinction in latitude, catchment properties, and temperature (range 0.2 -24.7 °C), the sampled lakes varied in nutrient content. Total organic carbon (TOC) in the lakes ranged from 0.6 to 116 mg l^-1^ (with a median of 13.3 mg l^-1^ in Norwegian and 9.4 mg l^-1^ in Swedish lakes), total phosphorus (TP) from 2 to 136 µg l^-1^ (median of 8.9 µg l^-1^ in Norwegian and 10 µg l^-1^ in Swedish lakes), and total nitrogen (TN) from 50 to 1280 µg l^-1^ (median of 271 µg l^-1^ in Norwegian and 360 µg l^-1^ in Swedish lakes). Additionally, we recorded pH levels ranging from 4.4 to 9.0 (median of 6.5 in Norwegian and 6.7 in Swedish lakes). In both datasets, TOC, TP, and TN correlated positively with each other and with turbidity, and negatively with latitude (Figure S2). The Norwegian lakes varied in size from 0.08 to 14.5 km^2^ with a median of 0.98 km^2^. In contrast, the Swedish lakes exhibited sizes ranging from 0.03 to 14 km^2^ with a median of 0.44 km^2^ (Table S1).

### Fungal contribution and alpha diversity in freshwater lakes

Utilizing high-throughput sequencing of the eukaryotic SSU rRNA gene, our study revealed that fungal taxa constituted a substantial portion of the overall diversity within some of the studied lakes, spanning a wide range from 0.18% to as much as 55% (Figure S3). Specifically, the Norwegian dataset displayed an average of 2.07% fungal reads, while the Swedish dataset exhibited an average of 5.41% fungal reads. These observations align with previously reported ranges for fungal relative read abundances in lakes ^7,14,34^ and ponds ^35^. These findings are also consistent with studies involving measurements of phospholipid-derived fatty acids in seston biomass of numerous lakes and kettle holes in northeast Germany, where fungi typically accounted for an average of 9.2 ± 5.2% of the biomass in the pelagic zone ^36^.

Our analysis, employing partial least squares (PLS) modeling, revealed that the relative abundance of fungi in relation to other eukaryotic SSU rRNA genes increased with catchment land cover classified as forests and decreasing pH values in the Swedish dataset while in the Norwegian dataset, these patterns were not clear (Figure 1). Noteworthy, the highest fungal relative abundances were found in dystrophic (high proportion of forests in the catchment) lakes. Dystrophic lakes are often characterized by anoxic bottom waters (hypolimnion) and high inputs of terrestrial organic matter giving them their distinctively dark, tea-stained appearance. This reflects that fungi, with larger genomes and higher enzymatic capabilities, can thrive best in lakes with a higher content of recalcitrant organic matter, which is similar to the reported for ectomycorrhizal fungi in terrestrial environments ^37^. Resulting low light conditions, and nutrient limitations, especially when it comes to nitrogen and phosphorus, restrict primary productivity in dystrophic lakes, tipping the balance towards heterotrophic eukaryotes such as fungi.

**Figure 1.**
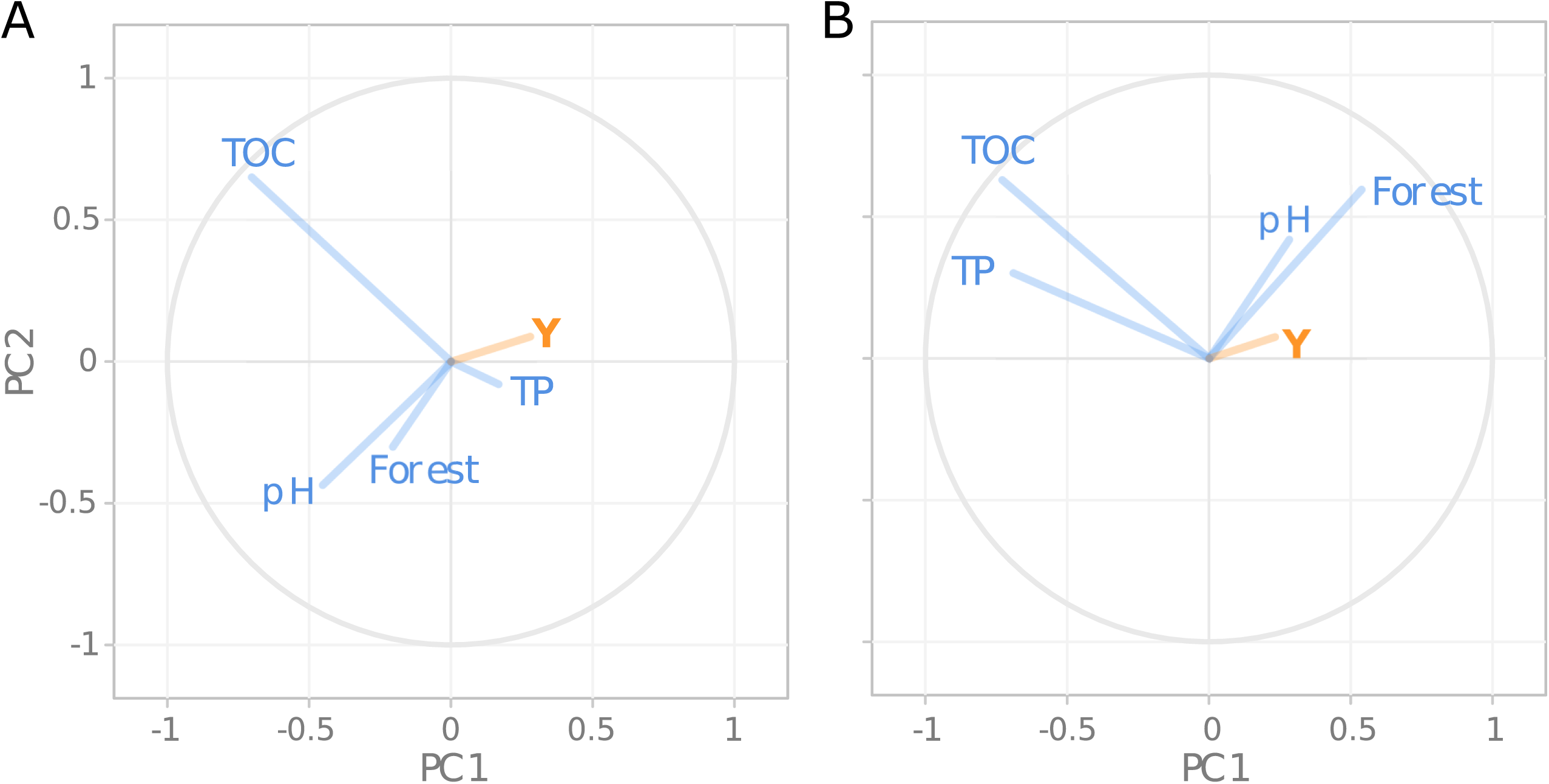
Plot representing the results from a partial least squares regression model that predicts fungal contribution (in orange) to the eukaryotic community from environmental data (in blue). Environmental variable arrows with the smallest angle (i.e. closest) to the eukaryotic arrow (‘Y’) have the highest correlation with fungal contribution. The models are based on data from 73 Norwegian (A) and 83 Swedish (B) lakes.

The fungus-specific sequencing libraries, covering the partial 5.8S gene and ITS2 region, yielded 631 and 1271 fungal operational taxonomic units (OTUs) in the Norwegian (comprising 51 lakes) and Swedish (comprising 84 lakes) datasets, respectively. After rarifying to 700 reads, observed OTU richness (ranging from 3 to 79) and evenness (Pielou’s evenness ranging from 0.02 to 0.94) varied widely across the datasets but could not be explained by any of the environmental properties as tested by correlation and PLS analysis. In the fungus-specific sequencing libraries, percentages of OTUs assigned to the kingdom *Fungi* varied from 44% for the Norwegian lakes to 59% for the Swedish ones, which is in line with previous results from surface waters in Iceland, where 54% of the reads were recovered as Fungi ^38^. A high proportion of OTUs were not assigned at the kingdom level (33 and 21% for Norwegian and Swedish datasets, respectively) or matched non-fungal organisms (23 and 20%), mainly members of the kingdoms *Alveolata* (phylum *Ciliophora*) and *Viridiplantae*, reflecting the relatively low specificity of the selected fungal primers (ITS3-Mix2 and ITS4-cwmix1/2). Similar results were obtained in a previous fungal DNA metabarcoding study, using the primers fITS7a and ITS4 to amplify the ITS2 region, on Scandinavian lakes ^14^. Comparing it to the eukaryotic data (SSU libraries), both *Alveolata* (*Ciliophora*) and *Viridiplantae* (*Chlorophyta*) were found to be prevalent and abundant (Figure S3).

A large portion of the fungal OTUs was unique to specific lakes, with 72 % (438) of Norwegian and 71% (549) of the Swedish OTUs occurring in only a single lake. An additional 19 % were observed in two or three lakes within the Norwegian and Swedish systems. The remaining 9.7 % of Norwegian (51) OTUs and 10.0 % of Swedish (93) OTUs were distributed across more than three lakes. These common OTUs often dominated the fungal community, with a combined relative abundance ranging from 0.3 % to 99 % per Norwegian lake and 0% to 100% per Swedish lake, averaging 54% and 71% across the Norwegian and Swedish datasets, respectively (Figure S4).

Within the Norwegian dataset, the most prevalent OTU, identified as *Hyaloscyphaceae* (*Ascomycota*), was found in 20 lakes, while the dominant OTU in the Swedish dataset was an unassigned *Rozellomycota* occurring in 49 lakes. The presence of numerous OTUs restricted to specific lakes, along with low prevalence among most OTUs (see Figure S5), suggests a high level of among-lake richness. This was further substantiated by species accumulation curves (SAC) (see Figure S6), indicating that the OTU pool remains undersampled across the broad environmental gradients sampled, despite the sequencing efforts that detected 631 and 1271 OTUs in the rarefied Norwegian and Swedish datasets, respectively. Most likely, the limited specificity of ITS primers discussed above has decreased the effective sequencing depth for recovering the full fungal diversity of the studied samples. In addition, it is important to note that these results, based on denoised data designed to remove artificial diversity introduced during sequencing, may still contain incorrect sequences, potentially inflating richness and distorting community composition ^39,40^.

### Deterministic versus stochastic fungal community assembly

A first visual inspection of beta diversity using nonparametric multidimensional scaling (NMDS) did not reveal any clear latitudinal pattern for both datasets (Figure S7). Redundancy analysis (RDA) and fitting environmental variables to the ordinations as well as partial mantel tests suggested that fungal diversity was only partly driven by the measured environmental variables. The variance explained by RDA models was around 5% using different sets of environmental variables. Partial mantel tests were inconclusive since in the Norwegian dataset geographic distance and in the Swedish dataset environmental properties explained around 20% of the variability (see Table S2 for details). Such results provide indications that besides profound differences in explanatory variables, deterministic processes such as environmental sorting and dispersal can partially overwrite random ecological drift (stochastic processes).

To dissect the interplay between stochastic and deterministic processes shaping fungal communities, we conducted Monte Carlo simulations to analyze incidence-based co-occurrence patterns ^41^ and Raup-Crick-based beta-diversity (*β_RC_*)^26^ of the OTUs within the study lakes. Our findings revealed Standard Effective Sizes that did not consistently significantly deviate from random within the Swedish dataset across the co-occurrence statistics such as the C-score - *cscore*, count of checkerboard species pairs - *checker*, and the number of species combinations – *combo*, when employing different randomization approaches considering fixed, equiprobable, or proportional constraints for rows and columns (Table 1). In the Norwegian dataset, determinism in cooccurrence patterns was widely observed (Table 1), and *β_RC_* values <-0.95 and >0.95 were overrepresented (Figure 2), pointing to competitively structured communities. Values of *β_RC_* varied widely but showed an increased number of values <-0.95 and >0.95 in the Swedish dataset, although mean values did not deviate from the null expectations (Figure 2). Consequently, these findings suggest that ecological drift plays a pivotal role in the assembly of fungal communities ^25^ across Scandinavia although deterministic processes like environmental filtering and species interactions ^25,42^, as well as biogeographic and evolutionary history ^43,44^ can be important but vary over space and time ^20^.

**Figure 2.**
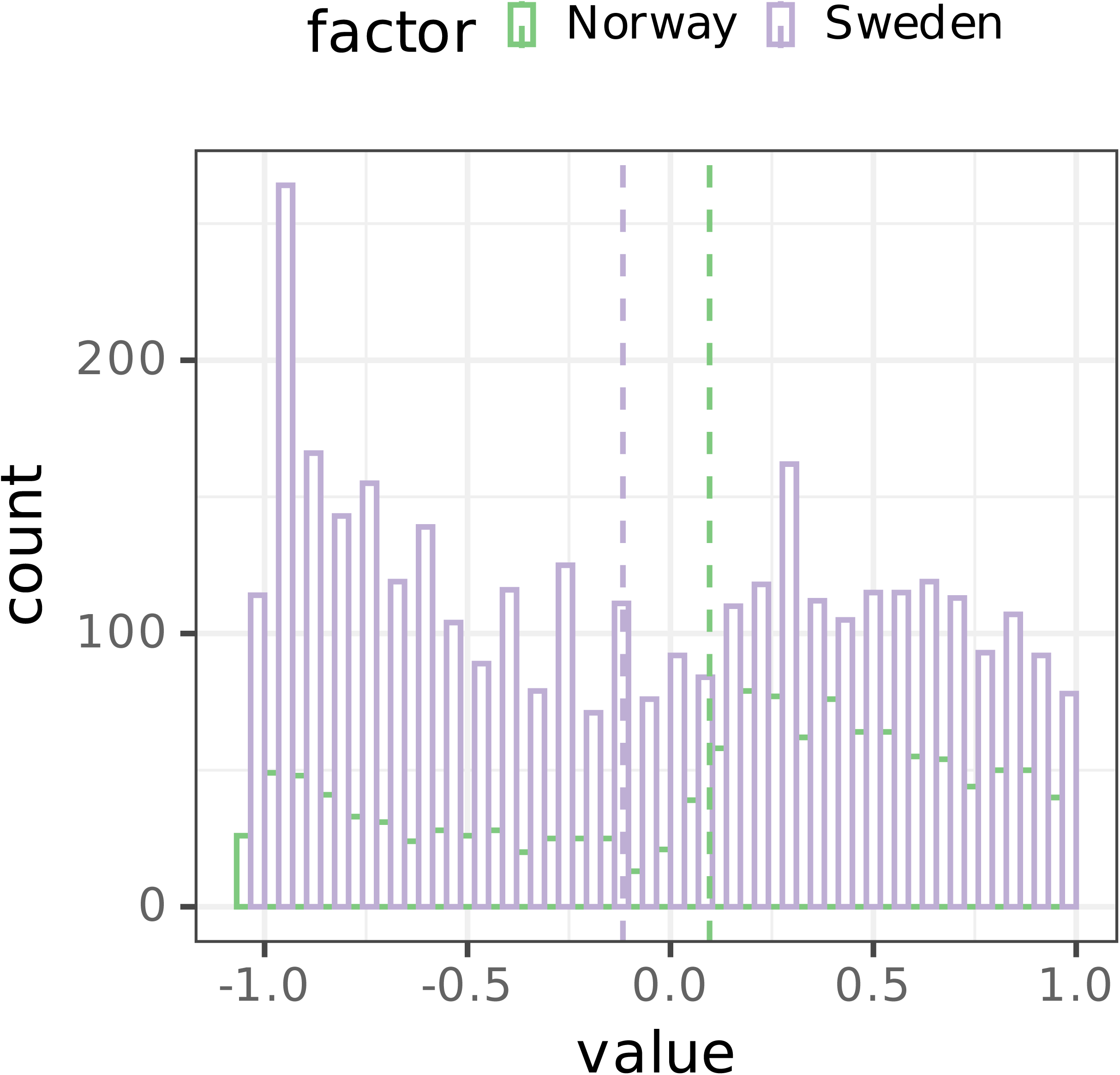
Variation of incidence-based (Raup-Crick) beta-diversity (β_RC_) in the fungal communities from the Norwegian (green; 61 lakes) and Swedish (blue; 83 lakes) datasets. Dashed line refers to the mean values of the respective lake dataset.

**Table 1.**
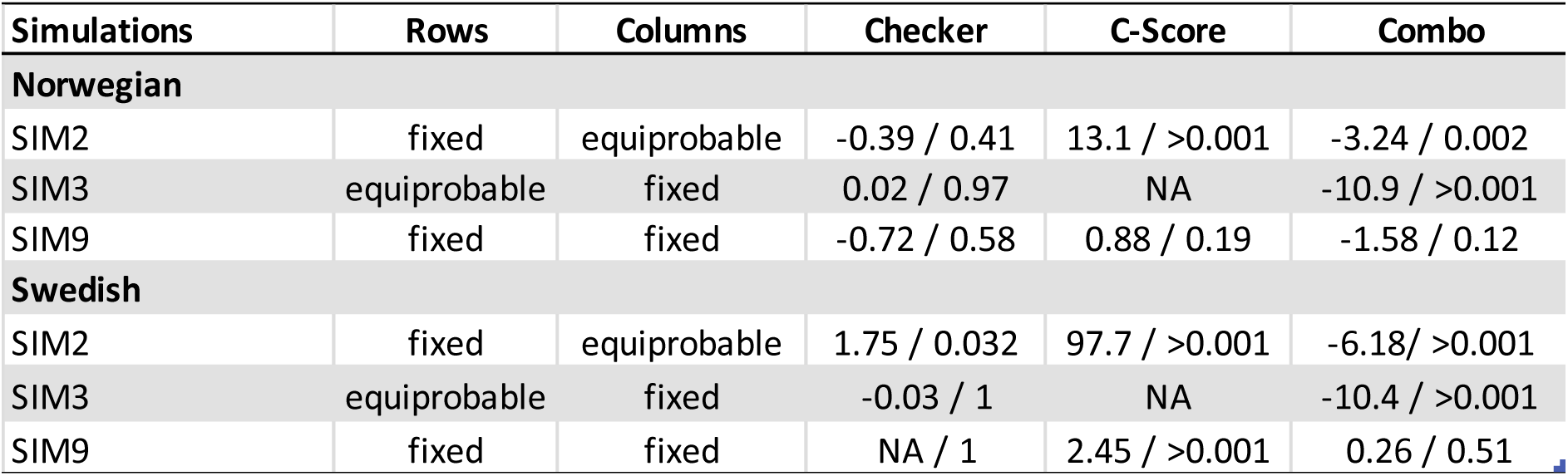
Results from incidence-based EcoSimR ^41^. Utilizing Monte Carlo randomizations z-scores for various co-occurrence indices including C-score ^79^, the number of checkerboard species pairs ^42^ (Checker), and the number of species combinations ^23^ (Combo), were estimated for the Norwegian and Swedish fungal lake amplicon variants (OTUs). Fixed rows-fixed columns and fixed rows-equiprobable columns simulations were utilized to minimize Type I and Type II errors ^25^.

To further resolve and estimate the relative importance of these deterministic assembly processes, we applied the quantitative process estimate (QPE) framework ^22^. The results showed some differences between the two datasets (Figure 3). In both the Norwegian and Swedish datasets, ecological drift was the dominant assembly process based on pairwise comparisons (*β_NTI_* <2, and *-0.95> β_RC_*values <0.95; 86.18 and 83.64% of pairwise comparisons in the Norwegian and Swedish datasets, respectively). Homogenizing dispersal or mass effects (*β_NTI_* <2, and *β_RC_* values <-0.95; 9.18 and 12.83% of pairwise comparisons in the Norwegian and Swedish datasets, respectively) were followed by dispersal limitation or historical contingency (*β_NTI_* <2, and *β_RC_* values >0.95; 4.64 and 3.53% of pairwise comparisons in the Norwegian and Swedish datasets, respectively), while homogeneous selection (*β_NTI_* >2) was not identified. The low proportions of homogeneous selection (0%) and dispersal limitations (3.5–4.7%) confirm their minor role in fungal assembly as inferred from partial mantel tests (Table S2). These results suggest that lake fungal communities are primarily structured by stochastic processes and that predictive models on fungal alpha and beta diversity, as shown in our efforts, are hard if not impossible to obtain.

**Figure 3.**
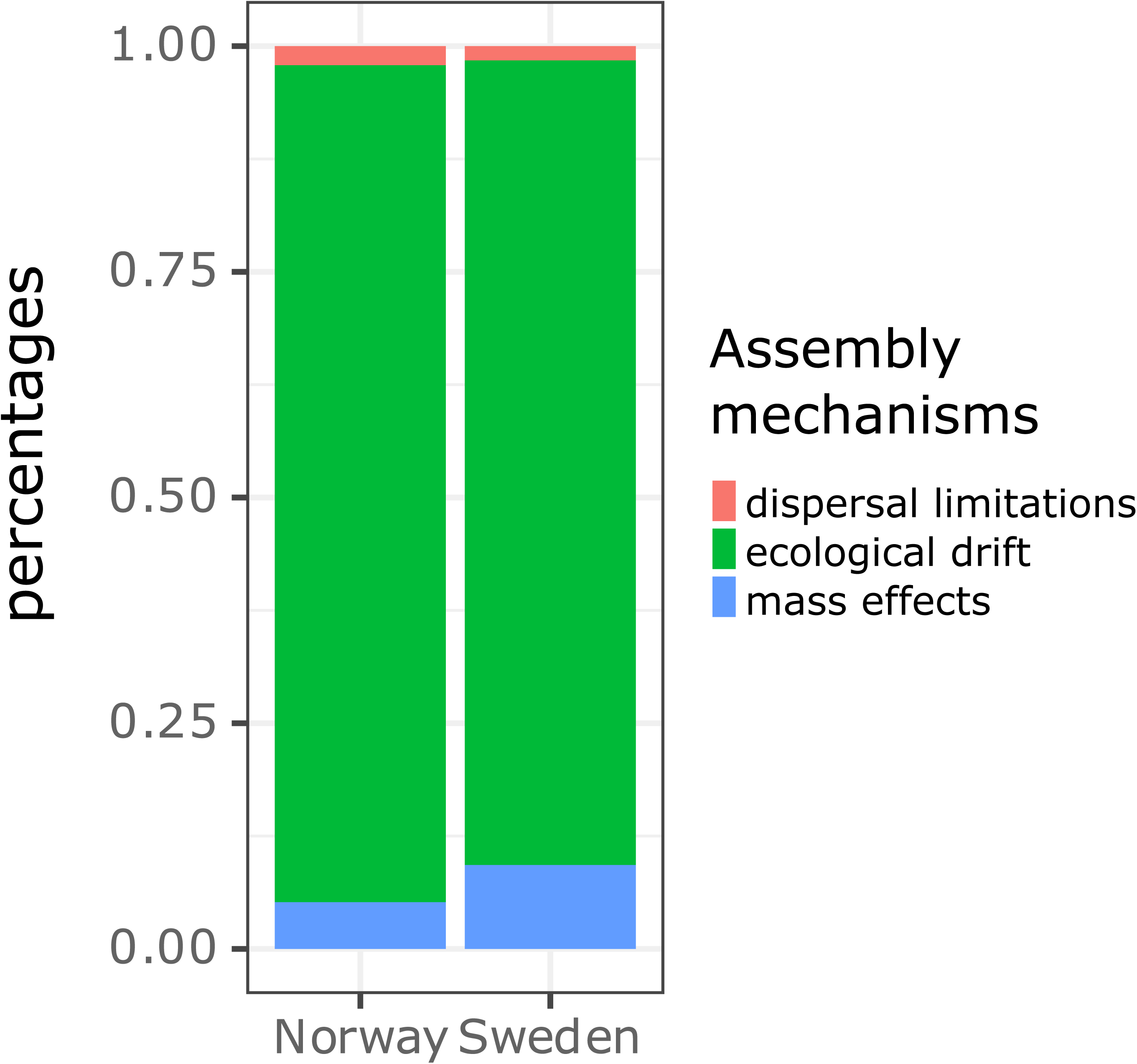
Relative importance of different community assembly processes expressed as the proportion of community pairs assembled either by species-sorting (variable or homogeneous selection), dispersal limitation or historical contingency, homogenizing dispersal or drift for fungal communities in the Norwegian and Swedish lakes.

### Taxonomical distribution of fungi across Scandinavian freshwater lakes

The most abundant fungal phyla across the Norwegian lakes were *Ascomycota* (50% of the reads), *Rozellomycota* (19%), *Basidiomycota* (5.3%), *Chytridiomycota* (2.1%), *Aphelidiomycota* (1.9%), and *Glomeromycota* (1.7%). In Swedish lakes, the dominant phyla were *Rozellomycota* (29%), *Ascomycota* (28%), *Basidiomycota* (21%), *Chytridiomycota* (16.5%), and *Mortierellomycota* (0.2%).

The majority of reads of both datasets were not assigned at the genus level, with percentages of unidentified reads per sample ranging from 3 to 100% (Figure S8). The low assignment percentage can be partially due to the conservative method used for the taxonomical assignment by the lowest common ancestor analysis (LCA), as well as the poor representation of the aquatic (freshwater and marine) fungi in the UNITE reference database. Previous studies have identified the aquatic ecosystems as priority targets for the recovery of novel lineages within the early diverging fungi affiliated to phyla such as *Rozellomycota*, *Chytridiomycota,* and *Aphelidiomycota* ^2^, which were abundant in our survey, and especially in the Swedish dataset.

Regarding the limited fraction of OTUs identified at the genus level, the most common genera in the Norwegian dataset were taxa associated with plants (both saprobic and pathogenic fungi) such as *Ascocoryne, Ramularia, Naevala, Oidiodendron,* and *Lemonniera*, as well as the lichenicolous genus *Epithamnolia*. *Lemonniera* is a known aquatic genus inside *Leotiomycetes*^45^. In contrast, the most abundant genera in the Swedish dataset were yeasts belonging to the basidiomycete orders Tremellales (*Naganishia, Vishniacozyma,* and *Filobasidium*; which were formerly referred to the genus *Cryptococcus*) and *Sporidiobolales* (*Rhodotorula*), and the ascomycete order *Dothideales* (*Aureobasidium*). Previous studies have reported the genera *Cryptococcus*, *Rhodotorula,* and *Aureobasidium* in freshwater systems^2,6,14,46^. In addition, the ubiquitous mold genus *Alternaria* was also abundant in the Swedish dataset.

The higher number of genera identified in the Swedish (284 genera) compared to the Norwegian (159 genera) dataset is most likely due to the wider latitudinal range sampled. From the total of 355 genera identified in both datasets, 25% (89 genera) of these were shared among the Norwegian and Swedish datasets (Figure S9). Some of these overlapping genera have been identified in previous lake studies: *Aureobasidium, Candida*, *Chytridium*, *Rhodotorula*, *Trechispora,* and *Rozella* ^14,35,47^.

Our results on the fungal taxonomical composition of Scandinavian lakes support previous studies^14,48^, which reported that fungal communities of freshwater systems contain resident taxa, those active members of the community with a clear aquatic origin, as well as transient taxa, with a terrestrial origin, whose fungal structures have been washed into aquatic habitats^49^. For this comparison of Swedish and Norwegian data, the seasonality has to be taken into account. The Norwegian water samples were collected when the autumn leaf fall took place and thus fungi linked to plant material degradation were prominently detected, while the August samples from the Swedish lakes instead reflected planktonic fungi that are linked to the autotrophic food-web, like yeasts and parasites (*Rozellomycota*).

Redundancy analysis indicated that the measured environmental lake properties could only explain a minor fraction of the taxonomic shifts at the phylum and genus levels (Table S2). Using generalized linear latent models (gllvms) on individual genera revealed some strong relationships with lake properties such as nutrient status (i.e. total phosphorus) and total organic carbon concentrations as well as landscape features (% of forest in the catchment) (Figure 4, supplementary Figures S10-11). For example, *Anguillospora*, a genus previously described to be common in freshwaters, was predicted to increase in contribution in lakes with high nutrient content while being less common in forested lakes with high TOC concentrations. *Lemmoniera* is another common freshwater fungal genus that was positively correlated with nutrient status (i.e. TP) while no relationships with the other lake properties could be identified. As such the gllvms expand the knowledge about the ecological range of key freshwater fungal genera and provide insights into the ecological drivers of individual genera. This information is essential for species distribution modeling to better comprehend the depths of fungal biodiversity change, reliably anticipate these changes, and inform conservation actions ^50^.

**Figure 4.**
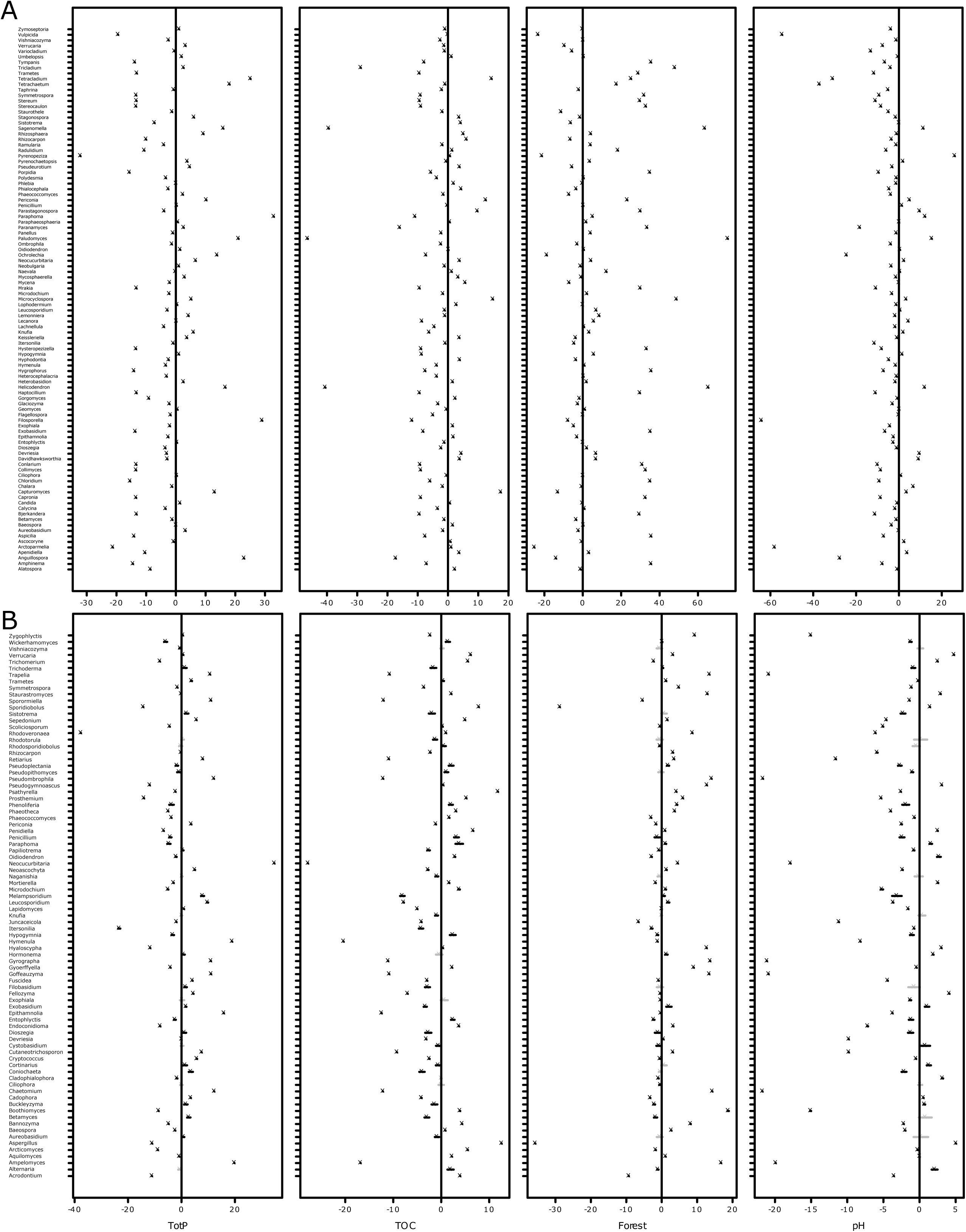
Results from a general linear latent model (gllvm) on fungal genera from Norwegian (A) and Swedish (B) lakes. The estimated coefficients for predictors and their confidence intervals (TP – total phosphorus, TOC – total organic carbon, Forest – forest coverage in the catchment, and pH), allow to study the nature of effects of environmental variables on fungal genera. Point estimates (ticks) for coefficients of the environmental variables and their 95% confidence intervals (lines) are given, with those colored in black denoting intervals not containing zero. Evaluation statistics of the models are shown in Figures S10 and S11.

## CONCLUSION

Our observations do not fully support the predictions of the size-dispersal hypothesis, which posits that small organisms, such as most fungi, are primarily structured deterministically due to their susceptibility to species sorting rather than dispersal limitations. This is attributed to the fact that organisms at the µm-scale possess a remarkable capacity for dispersal ^51–53^. According to the species sorting perspective, dispersal is vital for species to adjust to variations in local conditions, ultimately leading to deterministic processes overriding ecological drift ^54^. However, the excessive dispersal (homogenizing dispersal; *β_NTI_* <2, and *β_RC_*values <-0.95) that transfers substantial numbers of organisms across ecosystem boundaries—commonly referred to as "mass effects"— attenuates the impact of local environmental conditions ^55^, potentially distorting selection processes, as reflected in all pair-wise comparisons having a *β_NTI_* <2. Furthermore, variations in species sorting mechanisms, as indicated by the results from the gllvm, across fungal taxa can introduce additional stochasticity.

Another important aspect promoting overall stochasticity (*β_NTI_* <2, and *-0.95> β_RC_* values <0.95) in addition to ecological drift, is the variability in dispersal mechanisms among different fungal taxa ^56,57^, contributing to the absence of a universal dispersal process across the surveyed landscapes ^57^. This is reflected in most pairwise sample comparisons having *β_RC_*values ranging between 0.95 and -0.95. Not determined, recent diversification events and any historical impediments to gene flow ^58^ can also contribute to stochasticity, thus presenting a major challenge in untangling the intricacies of community assembly processes ^59–62^, and thus modeling alpha and beta diversity dynamics across climate zones.

## METHODS

### Sampling Norwegian lakes and measurements of environmental state variables

In the autumn (October – November) of 2019, we gathered surface water samples from 73 lakes in south-eastern Norway. These lakes were specifically selected as a subset of those monitored in a comprehensive 1000-lake survey across Norway, conducted by the Norwegian Institute of Water Research (NIVA). This synoptic survey aimed to replicate earlier campaigns carried out in 1986 and 1995 ^63^, and covered a broad spectrum of water quality properties, including dissolved organic matter (DOM) and nutrients ^64^, as well as catchment properties and land use ^65^. Sampling took place in late autumn, coinciding with or following lake overturn. Samples were collected approximately 4 meters from the lake’s shoreline using a sampling rod equipped with a sampling beaker. Whenever feasible, we utilized the lake’s outlet as the sampling point. A composite water sample of approximately 2 liters was collected in a sampling bucket with minimal physical disturbance. For DNA analyses, water samples (from 0.2 to 1.2 liters) were filtered through 0.2-μm pore size Sterivex cartridges (Millipore) in duplicate, using sterile syringes. These filters were then immediately frozen in liquid nitrogen on-site until further processing. Upon sample collection, we immediately measured water temperature (T), pH, and electrical conductivity (EC). Additionally, 50 milliliters of unfiltered water were collected from each site and stored at 10°C in polypropylene tubes. Upon arrival at the laboratory, these samples were frozen at -20°C until analysis for TOC, TN, and TP at the University of Oslo. TOC was determined by infrared CO_2_ detection following catalytic high-temperature combustion using a Shimadzu TOC-VWP analyzer. TN was quantified by detecting nitrogen monoxide through chemiluminescence with a TNM-1 unit attached to the Shimadzu TOC-VWP analyzer. TP was measured as phosphate after wet oxidation with peroxodisulfate using an auto-analyzer.

### Sampling of Swedish lakes and measurements of environmental state variables

In August 2014, as part of a comprehensive national lake monitoring program (for more details see ^19^), we obtained samples from 119 Swedish lakes. Water samples for chemical analysis were acquired from a depth of 0.5 meters using a 0.5-meter long tube sampler of Ruttner-type design. For microbiological analysis, samples were collected from the entire epilimnion, typically extending between 2 and 8 meters in depth, employing a 2-meter long tube sampler. These epilimnion samples were combined in a large container, and subsequently, subsamples of 100 and 500 milliliters were filtered using a peristaltic pump through 142 mm Millipore polycarbonate filters with a 0.2-μm pore size, continuing until the filters became clogged. The filters were promptly stored at -80°C until further processing. Geographical data, including lake area and catchment area, were sourced from the Svenskt Vattenarkiv (Swedish Water Archive, SMHI, 2012) database. Information on catchment land use and cover, such as the percentages of forest, agriculture, and urban areas, were obtained from the Swedish Land Cover Database (Naturvårdsverket, 2014), which is a subset of the CORINE Land Cover data (European Environment Agency, 2014).

Water physicochemical data at the time of sampling, collected as part of the Swedish national lake monitoring program, are publicly accessible through the national data host (https://www.slu.se/en/departments/aquatic-sciences-assessment/data-host/). These physicochemical measurements adhered to international (ISO) or European (EN) standards.

### DNA extraction and eukaryotic sequencing of Norwegian lakes

Total DNA was extracted from the Sterivex cartridges using the DNeasy PowerWater Sterivex Kit (Qiagen, Germany). Eukaryotic SSU rRNA gene amplicons were sequenced following established protocols ^66^. In short, eukaryotic primers TAReuk454FWD1 5’-CCAGCASCYGCGGTAATTCC-3’ and V4 18S Next.Rev 5’-ACTTTCGTTCTTGATYRATGA-3’ ^67^ were utilized, equipped with Nextera adapters and indices. The pooled samples were subsequently sequenced at IMR sequencing facility (Dalhousie University, Halifax, Canada) using an Illumina MiSeq machine with a 2x300 bp chemistry, yielding sufficient high-quality sequences from all 73 Norwegian lake samples.

### DNA extraction and eukaryotic sequencing of Swedish lakes

DNA extraction was made from polycarbonate filters using the PowerSoil DNA Isolation Kit (MO BIO, USA), followed the method outlined by Juottonen and colleagues ^19^. For the preparation of eukaryotic libraries, a two-step PCR protocol was employed to minimize potential biases associated with PCR, such as chimera and heteroduplex formation ^68^. Additionally, this protocol allowed for the incorporation of barcodes, Illumina handles, and adapters into individual sample amplicons. In brief, approximately 1 ng of genomic DNA was added to duplicate PCR reactions containing dNTPs (0.2 mM), the eukaryotic primers 574*f (5′-ACACTCTTTCCCTACACGACGCTCTTCCGATCT-NNNN-CGGTAAYTCCAGCTCYV-3′) and 1132r (5′-AGACGTGTGCTCTTCCGATCT-CCGTCAATTHCTTYAART-3′) ^69^ at 0.25 µM, as well as Q5 High-Fidelity DNA Polymerase (New England Biolabs Inc; 0.4 unit) and PCR buffer (5 ×), in a final volume of 20 μl. The amplicon size ranged between 700-800 bp. The first PCR involved an initial denaturation step at 98 °C for 1 min, followed by 20 cycles consisting of 10 s at 98 °C, 30 s at 51 °C, 1 min at 72 °C, and a final elongation step of 2 min at 72 °C. Duplicate amplicons were subsequently pooled and purified using the Agencourt AMPure XP purification system. A second PCR step was performed, utilizing the same PCR reagents/concentrations except for primers (detailed below), for 15 (archaea) or 12 cycles (eukaryotes). This step included an initial denaturation at 98 °C for 30 s, followed by 10 s at 98 °C, 30 s at 66 °C, and 72 °C, with a final extension step of 2 min at 72 °C. The previous PCR product (1 μl) served as the template for this second PCR, which introduced barcodes and complete Thruplex adapters for Illumina sequencing, with forward primer 5’-AATGATACGGCGACCACCGAGATCTACAC-[i5 index]-ACACTCTTTCCCTACACGACG-3’ and reverse primer 5’CAAGCAGAAGACGGCATACGAGAT-[i7 index]-GTGACTGGAGTTCAGACGTGTGCTCTTCCGATCT-3’. Following a purification step with the Agencourt AMPure XP kit, quantification was conducted using a PicoGreen assay (Quant-iT PicoGreen, Invitrogen). Subsequently, SSU rRNA gene samples were pooled equimolarly, and sequencing was performed on a MiSeq instrument using 2x300 bp v3 chemistry and software version 2.6.1.1 at the Science for Life Laboratory in Uppsala, resulting in high-quality sequencing data from a total of 83 Swedish lake samples.

### Fungal sequencing

Fungal libraries were constructed employing a two-step PCR protocol with slight modifications. Genomic DNA ranging from 10-90 ng was added to duplicate PCRs containing dNTPs (0.2 mM), 0.4 units of Q5 polymerase, and PCR buffer (5×) in a final volume of 20 μl. Fungal-specific primers, including ITS3-Mix2 (5’-ACACTCTTTCCCTACACGACGCTCTTCCGATCT-NNNNNN-CAWCGATGAAGAACGCAG-3’) and an equimolar mixture of the reverse primers ITS4-cwmix1 (reverse; 5-GTGACTGGAGTTCAGACGTGTGCTCTTCCGATCT-NNNNNN-TCCTCCGCTTAYTGATATGC-3’) and ITS4-cwmix2 (reverse; 5’-GTGACTGGAGTTCAGACGTGTGCTCTTCCGATCT-NNNNNN-TCCTCCGCTTATTRATATGC-3’), were used. These primers were designed to include Illumina adaptors and fungus-specific sequences described by ^16,70^. The resulting amplicons were approximately 350-500 bp in length, encompassing approximately 130 bp of the 5.8S rRNA gene and the entire ITS2 region, as previously proposed by^38,71^.

The first PCR included an initial denaturation step of 98 °C for 1 min, followed by 30 cycles of 20 s at 98 °C, 30 s at 55 °C, 1.5 min at 72 °C, and a final extension of 7 min at 72 °C. High-quality sequences were obtained from 61 Norwegian lake samples and 83 Swedish lakes.

### Bioinformatic analyses

Raw sequence data from the eukaryotic and fungal sequences were trimmed of primers with CUTADAPT ^72^, and sequences without matching primers and fungal reads shorter than 200 bp were removed. Subsequently, each individual sequencing run (n = 4) was subjected to de-replication, denoising, and sequence pair assembly using the R package *dada2* ^73^. Quality score plots were manually inspected, and for Norwegian eukaryotic amplicons, forward and reverse reads were trimmed to 270 and 190 bp length, respectively, while the Swedish eukaryotic amplicons were trimmed to 190 and 120 bp length, respectively. Fungus-specific sequences were not truncated at this step. Additional quality filtering eliminated sequences with unassigned base pairs and reads with a single Phred score below 20. After dereplication, forward and reverse error models were established in *dada2*, employing a subset of sequences (10^7^ bases for eukaryotic and 10^9^ bases for fungal forward and reverse reads) from each sequencing set. Eukaryotic forward and reverse reads were concatenated while fungal paired-end reads were merged. Chimeric sequences were removed using the “removeBiomeraDenova” function within the *dada2* package, and amplicon sequence variants (ASVs) tables were obtained. Eukaryotic sequences were taxonomically assigned using the PR^2^ database ^74^.

Fungus-specific sequences underwent further processing: (i) extraction of the 5.8S rRNA gene and ITS2 region with ITSx, (ii – for ITS2) clustering of ASVs into OTUs at 97% similarity and removal of singletons using VSEARCH ^75^, (iii – for ITS2) curation of OTU tables with *lulu* ^76^, (iv - for 5.8S) dereplication and sorting using VSEARCH, (v – for both regions separately) taxonomic assignments of OTUs/ASVs against the UNITE database v. 8.3 (UNIT+INSD for eukaryotes) ^77^ using BLAST and lowest common ancestor analysis (LCA), and finally (vi) merging the two taxonomic assignments to generate the final taxonomy using R.

### Data visualization and statistical analyses

All statistical analyses and data visualization were conducted using R version 4.2.1, with multiple R packages employed (for specific details, refer to the deposited R code). Missing values in the metadata were approximated using multiple imputations via the MICE algorithm ^78^, which employed variables as predictors that met specific criteria: (i) inclusion in the complete data model, (ii) relevance to non-response, (iii) explanation of a significant portion of variance, and (iv) absence of excessive missing values within the incomplete cases’ subgroup. In cases of autocorrelation, a subset of variables was retained for downstream analysis. Partial least squares regression models were utilized to identify environmental variables predicting the contribution of fungal reads to the total read abundance in the eukaryotic dataset (SSU 18S rRNA gene).

### Relationships between environmental properties and fungal community composition

Before conducting alpha and beta diversity analyses, eukaryotic and fungal data (ASV and OTU matrices, respectively) were rarefied to the smallest sample size within the respective eukaryotic (3467 and 4430 reads) and fungal (700 reads) datasets using the “rrarefy” function in the *vegan* package. Alpha diversity measures, including ACE and Pielou’s indices, were computed based on the rarefied community data. Beta diversity matrices were calculated using Bray–Curtis dissimilarity on the rarefied and Hellinger-transformed community data. Distance-based redundancy analysis (dbRDA) was performed using the “capscale” function in *vegan* to identify environmental variables co-varying with fungal beta diversity. The significance of the dbRDA model was assessed through permutational analysis (using the “anova.cca” function), both for the overall model and individual factors (marginal effects). Additionally, environmental vectors and factors were fitted onto Bray–Curtis ordinations (i.e., NMDS) to obtain correlation coefficients and significance levels with beta diversity.

### Incidence-based co-occurrence patterns and beta-diversity (β_RC_)

To assess stochastic versus deterministic community assembly processes across the two Scandinavian lake gradients, we employed incidence-based (presence/absence) EcoSimR ^41^. This approach utilizes Monte Carlo randomizations to create “pseudo-communities”, enabling statistical comparisons with the patterns in the actual data matrix. Various co-occurrence indices, including C-score ^79^, the number of checkerboard species pairs ^42^, and the number of species combinations ^80^, were estimated. Fixed rows-fixed columns and fixed rows-equiprobable columns simulations were utilized to minimize Type I and Type II errors^25^.

Subsequently, we calculated the incidence-based (Raup-Crick) dissimilarity indices (*β_RC_*) to determine whether communities assemble stochastically or deterministically. We used the “raup_crick” function provided by ^28^. β_RC_ values not significantly different from 0 indicate stochastic assembly, while values close to −1 suggest deterministic processes favoring similar communities and those close to +1 favor dissimilar communities. To evaluate differences in β_RC_ values among different lake types, lakes were clustered based on environmental properties using k-means. The variables used in k-means clustering were chosen based on Spearman correlations and included the contribution of "Marsh"/"Wetlands", "Forest" to the catchment, total phosphorus, pH, temperature, and total organic carbon. Clustering quality was assessed using the Elbow, Silhouette, and Gap statistic methods ^81^.

### Quantitative process estimates (QPE)

To assess the relative contributions of potential species sorting, dispersal limitation, drift, and mass effects (referred to as “quantitative process estimates – QPE”), we adopted a two-step framework developed by ^22^. This approach requires that phylogenetic distances (PD) among taxa should reflect differences in the ecological niches they occupy, thereby carrying a phylogenetic signal. We tested the presence of phylogenetic signals using Mantel correlograms ^22^. Furthermore, the prerequisite of this framework was met as the RDAs and the fitting of environmental vectors to ordination results with respect to phylogenetic community composition showed no correspondence.

To initiate the QPE based on pairwise comparisons (among sites during each sampling occasion), our first objective was to determine the extent to which the observed *β_MNTD_* (β-mean-nearest-taxon-distance) deviated from the mean of the null distribution. We assessed its significance using the β-Nearest Taxon Index (*β_NTI_*), calculated as the difference between the observed *β_MNTD_* and the mean of the null distribution, expressed in units of standard deviations (SDs). When the observed βMNTD significantly exceeded (*β_NTI_* > 2) or fell below (*β_NTI_* < −2) the null expectation, it indicated that the community assembly was driven by variable or homogeneous selection, respectively. In cases where there was no significant deviation from the null expectation, the observed differences in phylogenetic community composition were attributed to dispersal limitation, homogenizing dispersal (mass effect), or random drift.

In the second step, we calculated the abundance-based beta-diversity using pairwise Bray-Curtis dissimilarity (*β_RCbray_*) ^22^. Communities that were not selected in the first step, thus not assembled by selection, were then characterized by (i) dispersal limitation coupled with drift if *β_RCbray_* > +0.95, (ii) homogenizing dispersal if *β_RCbray_*<−0.95, or (iii) random processes dominate (drift) if *β_RCbray_* fell in between −0.95 and +0.95. The fraction where *β_RCbray_* > +0.95 may indicate either “true” dispersal limitation or historical contingency, both resulting in more dissimilar communities than expected by chance. Throughout this manuscript, we use the term “dispersal limitation or historical contingency” to refer to this fraction. Differences in the QPE between the k-means clusters (as described earlier) were evaluated using Kruskal-Wallis tests.

To further investigate whether “true” dispersal limitation might have played a role, we conducted Mantel correlation analyses between geographic distances (Euclidean distances of geographical coordinates) and fungal community dissimilarities (*β_RCbray_*) using Spearman correlation and 999 permutations. A significant relationship between *β_RCbray_* and spatial distance would support the importance of dispersal limitation, while historical contingency would be deemed less significant.

Moreover, we performed Mantel correlation analyses between *β_RCbray_*and environmental dissimilarities (Euclidean distances) to explore whether community dissimilarities might be attributed to selection by environmental factors lacking a phylogenetic signal (“phylogenetically non-conserved selection”). Such factors might not have been detected in the first step but could be retained in the second step of the QPE analysis. To ensure that significant geographic distance effects were not confounded by spatially autocorrelated environmental variation, and vice versa, we conducted partial Mantel tests.

## Acknowledgements

We thank the Trend Lake national monitoring program in Sweden for taking extra samples for us during the regular sampling. The Norwegian lake sampling was carried out thanks to close cooperation between researchers from the Centre for Anthropocene Biogeochemistry (CBA, University of Oslo) and the NIVA. We want to in particular thank Jing Wei and Stina Drakare for their respective support in the Norwegian and Swedish sampling campaigns. Jing Wei and Pilar López Hernández performed the DNA extractions of the Norwegian and Swedish lake samples, respectively. The authors acknowledge all of the participants involved in the CBA-100 lakes survey, NIVA for providing depth data, and especially Per-Johan Færøvig and Berit Kaasa for their support with practicality issues in the labs. This research was supported by internal funds by the University of Oslo to A.E., funds from private foundations through UNIFOR to P.M-S, as well as funds from Lars Hiertas Minnesfond, Kapten Carl Stenholms Donationsfond, and Birgit och Birger Wåhlströms Minnesfond to H.N. H.J was funded by Carl Tryggers Foundation and N.V. by the Belmont Forum-BiodivERsA and the Research Council of Norway through ARCTIC-BIODIVER project. CW was funded by the German Research Foundation DFG WU890/2-1. M.J.L. thanks the Erasmus+ program from European Commission for supporting her traineeship at University of Oslo.

## Data availability

The raw demultiplexed sequence data (eukaryotic and fungal amplicon reads) generated in this study have been deposited in the Sequence Read Archive (SRA) under BioProject accession code PRJNA1118258.

## Code availability

The code used in this study is available at https://github.com/alper1976/fungal.

## Author Contributions

The research was conceptualized by P.M-S. and A.E. Fieldwork was conducted by L.F., N.V., E.W., A.E., and other members of the Centre of Biogeochemistry in the Anthropocene. Molecular analyses were performed by H.J., C.W., P.M-S., and M.J.L. The main analyses and visualization of the data were performed by P.M-S. and A.E. with support from H.J., E.W., and L.F. The first version of the manuscript was drafted by P.M-S. and A.E. All authors provided comments and were involved in writing the final version. Financial support for the project was acquired by P.M-S., H.N. and A.E. This manuscript has been proofread and edited with the help of chatGPT version 3.5.

